# Timing of locomotor recovery modulated by the *white* gene in *Drosophila*

**DOI:** 10.1101/032490

**Authors:** Chengfeng Xiao, R Meldrum Robertson

## Abstract

Locomotor recovery from anoxia follows the restoration of disordered ion distributions and neuronal excitability. The time taken for locomotor recovery after 30 s anoxia (around 10 min) is longer than the time for the propagation of action potentials to be restored (less than 1 min) in *Drosophila* wild-type. We report here that the *white* (*w*) gene modulates the timing of locomotor recovery. Wild-type flies displayed fast and consistent recovery of locomotion from anoxia, whereas mutants of *w* showed significantly delayed recovery. Genetic analysis including serial backcrossing revealed a strong association between the *w* locus and the timing of locomotor recovery, and haplo-insufficient function of *w*^+^ in promoting fast locomotor recovery. The locomotor recovery phenotype was independent of classic eye pigmentation, although both are associated with the *w* gene. Introducing up to four copies of mini-*white* (m*w*^+^) into w1118 was insufficient to promote fast and consistent locomotor recovery. However, flies carrying *w*^+^ duplicated to Y chromosome showed wild-type-like fast locomotor recovery. Furthermore, RNAi knockdown of *w* in neurons but not glia delayed locomotor recovery, and specifically, knockdown of *w* in putative subsets of serotonin neurons was sufficient to delay the locomotor recovery. These data reveal an additional role for *w* in modulating the timing of locomotor recovery from anoxia.

## Introduction

Adult *Drosophila* enter a coma-like state during exposure to anoxia. The changes due to anoxia in *Drosophila* photoreceptors as well as the mammalian central nervous system (CNS) include the occurrence of anoxic depolarization, disordered ion distributions and the loss of neuronal excitability (Hansen 1985; Agam *et al*. 2000; Takano *et al*. 2007). Once returned to normoxia the timing of recovery varies depending on specific processes. Recovery of neural K^+^ distribution after a <30s anoxia starts immediately and extracellular K^+^ concentration is almost back to normal within 30s (Armstrong *et al*. 2011). Propagation of action potentials recovers within 1 min whereas complex motor activity, such as continuous walking, takes longer (around 10 min) to recover (Krishnan *et al*. 1997). These observations indicate that the recovery of locomotor coordination involves not only the re-establishment of intercellular communication between neurons, and from neuron to muscle, but also the return of appropriate synaptic strength and coordination. Thus, the factors influencing ion channels, membrane fluidity, and vesicle recycling, together with the coordination of a variety of modulatory signaling cascades, could have considerable impact on locomotor recovery. The recovery process is complicated and so far little of the cellular or molecular basis regulating the timing of locomotor recovery from anoxia is understood.

The *white* (*w*) gene, discovered by Thomas Hunt Morgan in 1910 (Morgan 1910), encodes a subunit of a heterodimer transmembrane protein that functions as an ATP-binding cassette transporter (O’Hare *et al*. 1984; Hazelrigg 1987). The White protein is involved in the uptake of precursors into granules for the synthesis of pteridine (red) and ommochrome (brown) pigments (O’Hare *et al*. 1984; Dreesen *et al*. 1988; Tearle *et al*. 1989). In the Malpighian tubule, the *w*^+^ transport system is associated with the accumulation of intracellular 3-hydroxykynurenine and cyclic guanosine monophosphate (cGMP) in the granules/vesicles (Sullivan and Sullivan 1975; Evans *et al*. 2008). The *w*^+^ transport system is also proposed to transport the pigment precursors, guanine and tryptophan, crossing cytoplasmic membranes to contribute to the yet unknown metabolic routes for pigment synthesis (Green 1949; Sullivan *et al*. 1979; Sullivan *et al*. 1980; Ferré *et al*. 1986). Many other substrates, including biogenic amines (dopamine, serotonin and histamine), metabolic intermediates and small molecules are thought to be transported by this system (Anaka *et al*. 2008). Levels of dopamine and serotonin, important modulators of synaptic transmission, are reduced in the heads of *w* mutants relative to the wild-type (Borycz *et al*. 2008). Mutations at the *w* locus, including complete lack of transcription in null mutants, altered amounts of transcript, and mis-location to neighboring heterochromatin, are associated with impaired pigment transport and deposition in the compound eyes (Nolte 1950; Pirrotta and Bröckl 1984). In addition to the function for eye pigmentation, extra-retinal functions of *w* have been observed (Zhang and Odenwald 1995; Campbell and Nash 2001). In the context of locomotor recovery from anoxia, such a *w*^+^ transport system could have critical housekeeping functions in building up vesicle stores, promoting synchronous release and recycling of small molecules and facilitating fast recovery of ion disorders and neural transmission.

By characterizing the dynamics of locomotion before, during and after a transient anoxia, we observed that mutant w1118 flies took longer to recover locomotion compared with wild-type Canton-S (CS). The w1118 strain carries a null mutant allele of *w* in the X chromosome, and is heavily used as genetic background for generating transgenic lines. Both w1118 and CS flies are common controls for genetic study in *Drosophila*. In this report we provide evidence of the association between the *w* allele and locomotor recovery. Our data, collected using multiple approaches including serial backcrossing, RNAi knockdown and *w*^+^ duplication to Y chromosome, strongly support a conclusion that *w*^+^ modulates the timing of locomotor recovery from anoxia.

## Materials and Methods

### Fly strains

Fly strains used for the experiments were CS (#1, Bloomington stock center), w1118 (L. Seroude laboratory), other wild-types including Hikone-AS (#3), Amherst-3 (#4265), Florida-9 (#2374), and mutants including *w^1^* (#145), *w^a^* (#148) and *w^cf^* (#4450). The other fly strains and their sources were: duplication line Raf^11^/FM6, l(1)FMa^1^/Dp(1;Y)B^S^*w*^+^*y*+ (#5733), elav-Gal4 (#8765). repo-Gal4 (#7415), UAS-w-RNAi (1) (#31088) and UAS-w-RNAi (2) (#35573), *yw*ΔΔ1;cn,bw (L. Seroude laboratory), Janelia UAS lines pJFRC2 (#32189), pJFRC5 (#32192) and pJFRC7 (#32194) (Pfeiffer *et al*. 2010), UAS-hsp70 (Xiao *et al*. 2007), UAS-Httex1-Qn-eGFP (n = 47 or 103) (Zhang *et al*. 2010); UAS-hsp26 and UAS-hsp27 (Wang *et al*. 2004), UAS-w-eYFP (Evans *et al*. 2008), R50E07-, R50H05- and R50E11-Gal4 (Pfeiffer *et al*. 2008). *w*^+^; *cn,bw* were generated according to standard genetic methods in *Drosophila* (Greenspan 2004). The pJFRC2-7 flies for the *mw*^+^ test were backcrossed into w1118 background according to previous protocols (Garfinkel *et al*. 2004). UAS-*w*-RNAi flies were backcrossed to CS background by a two-step procedure. Male RNAi flies were first crossed into *w*^+^;;TM3/TM6, and male progeny (*w*^+^/y;;UAS-*w*-RNAi/TM3) were backcrossed into *w*^+^;;TM3/TM6 for establishing *w*^+^;;UAS-*w*-RNAi stock.

Flies were maintained on standard cornmeal medium at room temperature (21-23°C) in 12h/12h of light/dark illumination with light on at 7am and off at 7pm. Adult flies were collected within 2 days after emergence, transferred into fresh food vials and subjected to locomotor assay at least 3 days after collection. A period of 3 days free of nitrogen exposure was guaranteed before experiments. Experiments were performed during the daytime between 10 am and 4 pm to avoid the morning and evening locomotor peaks (Stoleru *et al*. 2004).

### An apparatus for locomotor assay

An apparatus for video capturing was used for the locomotor assay of multiple flies at the same time (Xiao and Robertson 2015). Briefly, the circular arenas (1.27cm diameter) were drilled in a 0.3 cm-thick Plexiglas plate. The 0.3 cm thickness allowed flies to turn around but suppressed the vertical walk, thus the captured 2D videos represented locomotion constrained to the horizontal plane. The bottom surface of the arenas was covered with chromatography paper (Cat# 05-714-4, Fisher Scientific) for air circulation. The top was covered with another sliding and slightly larger Plexiglas sheet with holes (0.3 cm diameter) close to one end for fly loading. The plate was then positioned in a larger container designed for nitrogen exposure. The plate was illuminated with a white light box (Logan portaview slide/transparency viewer). White cardboard screens were used for light reflection to improve illumination from the sides, and to shield the arenas from light in the room. Locomotor activities of flies were recorded with a video camera (Logitech Webcam C905) and its associated software. The grey-scale videos, formatted as Windows Media Video (WMV) at a frame rate of 15fps and a resolution of 1600×1200 pixels, were taken and stored for post analysis. Experimental settings, including illumination, light reflection, and camera configuration remained consistent from experiment to experiment.

### Analysis of locomotor recovery from anoxia

Locomotor activities before, during and after a short period of anoxia (pure nitrogen exposure at a speed of 10L/min) were analyzed. Flies were exposed to a 30s anoxia after 5 min of pre-anoxia locomotion and then observed for 1h during the recovery under normoxia. A slow air flow (2 L/min) was provided throughout the experiment, except for the period of 30s nitrogen exposure, to remove the effect of dead space. A prior acclimatization period (5 min) was allotted before the experiment to avoid initial elevated activities (Liu *et al*. 2007). We chose a 30s nitrogen pulse because male flies usually entered anoxic coma around 10-15s from the onset of anoxia in our experimental settings, and 30s exposure was sufficient to knockdown all flies without exception. During recovery most of the mutants started to walk continuously within 30 min whereas a few mutants required a longer time to walk. We therefore recorded 1h recovery time after anoxia. Preliminary experiments showed that, although the flies started to walk within an hour, normal locomotor activities were not completely recovered to the non-exposed control level in w1118 flies. After one day, recovery from the 30s anoxia is likely complete. Thus a period of 3-day free of anoxic exposure was guaranteed before experiments to remove or minimize the anesthetization effect during fly collection. Experimental flies were no more than 9 days old.

### Fly tracking

Fly tracking was performed according to the protocol reported previously (Xiao and Robertson 2015). Briefly, a script was developed to track fly position using free software Open Computer Vision 2.0 (0penCV2.0) operating in the environment of Microsoft Visual C++ 2008 Express. The path length per second was calculated using the positional information once every 0.2s. The main scripting procedures included, (1) learning a specific background from multiple frames of a video, (2) comparing the difference between each frame and the background, (3) computing the center of mass of each fly, and (4) calculating path length per second. A metric ruler was placed in the camera view alongside the arenas for pixel-mm conversion.

### Evaluation of time to locomotor recovery

Preliminary data showed that wild-type flies were not ready to walk continuously until around 10 min after 30s anoxic exposure. There were variations for the time to start locomotion from fly to fly and from strain to strain. A threshold criterion was established to evaluate the time required to walk continuously, which we termed as “locomotor recovery”. Definition of a threshold was based on 5 min locomotion before anoxia. In a pre-test with w1118 flies, the 25th percentile of 300 values of path length per second (equivalent to 5 min locomotion) was around 0.3 cm. We thus used 0.3 cm of path length per second as the threshold for determining locomotor recovery. Additionally, to avoid taking into account early sporadic events (i.e. wing closure or body roll-over) that could result in occasionally above-threshold path lengths, a criterion of at least 10 instances of path length per second at or above threshold (0.3 cm/s) within 60 consecutive seconds was set as the standard for recovered locomotion. The time to locomotor recovery was measured as the earliest time to meet this criterion.

### Immunohistochemistry

Experimental procedures were based on two protocols (Wu and Luo 2006; Jenett *et al*. 2012) with minor modification. Briefly, dissected brain tissues were fixed in freshly prepared 4% paraformaldehyde overnight. After three washes with PAT (PBS with 0.5% BSA and 0.5% Triton X-100), the tissues were incubated with blocking buffer (5% goat serum in PAT) at room temperature for 1h. The tissues were then incubated with primary mouse anti-GFP supernatant (12A6, DSHB, Iowa City, IA) at 1:20 in blocking buffer for 1-2 days. Following three washes the tissues were then incubated with secondary antibodies Alexa Fluor 488 conjugated goat anti-mouse IgG (115-545-003, Jackson ImmunoResearch, West Grove, PA) at 1:500. Secondary incubation was performed in the dark for 1-4 days. After three washes tissues were suspended in 200 μl SlowFade Gold antifade reagent (S36938, Life Technologies, Burlington, ON) and mounted on slides for microscopy. An image stack was taken using a Carl Zeiss LSM 710NL0 laser scanning confocal/multiphoton microscope and processed with associated software Zen 2009 (Carl Zeiss).

### Statistics

Because part of the data showed non-Gaussian distribution, non-parametric tests (Man-Whitney test or Kruskal-Wallis test) were performed to compare the difference of medians between two or multiple groups. TR was evaluated from eight flies (n = 8) for each test. More than three duplications of tests were conducted. Specific statistical methods are provided in the text or figure legends where applicable.

## Results

### Delayed locomotor recovery from anoxia in w1118 strain

During recovery from anoxia flies performed sequential actions in the following order: rhythmic leg twitching, wing closure, rolling over, standing upright, antennal grooming and walking. Most of the recovery time is taken from the return to normoxia to rhythmical twitching of legs, and from standing upright to walking. Wild-type flies do not walk continuously until around 10 min after a 30s anoxic exposure. We examined locomotor activity in adult flies by computing path length per second before and after an anoxia using an assay described previously (Xiao and Robertson 2015). Using path length information we evaluated the time required for the locomotor activity to be recovered to a threshold level, which we termed time to recovery (TR) (see Materials and Methods). CS and w1118 male flies walked continuously in small circular arenas (1.27cm diameter) under normoxia. A 30s anoxia was sufficient to induce coma in all flies after which locomotor activity gradually recovered (Figure 1A and B). CS male flies recovered with a median of 530.0 s (interquartile range (IQR) 461.8 - 553.0 s, n = 8), whereas locomotor activity of w1118 males recovered at a median of 1037.0 s (IQR 755.3 - 1797.0 s, n = 8). The w1118 males displayed longer TR than CS males (*P* < 0.05, *t* test with Welch’s correction), and larger variance of TR than CS males (*P* < 0.0001, F test) (Figure 1C).

**Figure 1.**
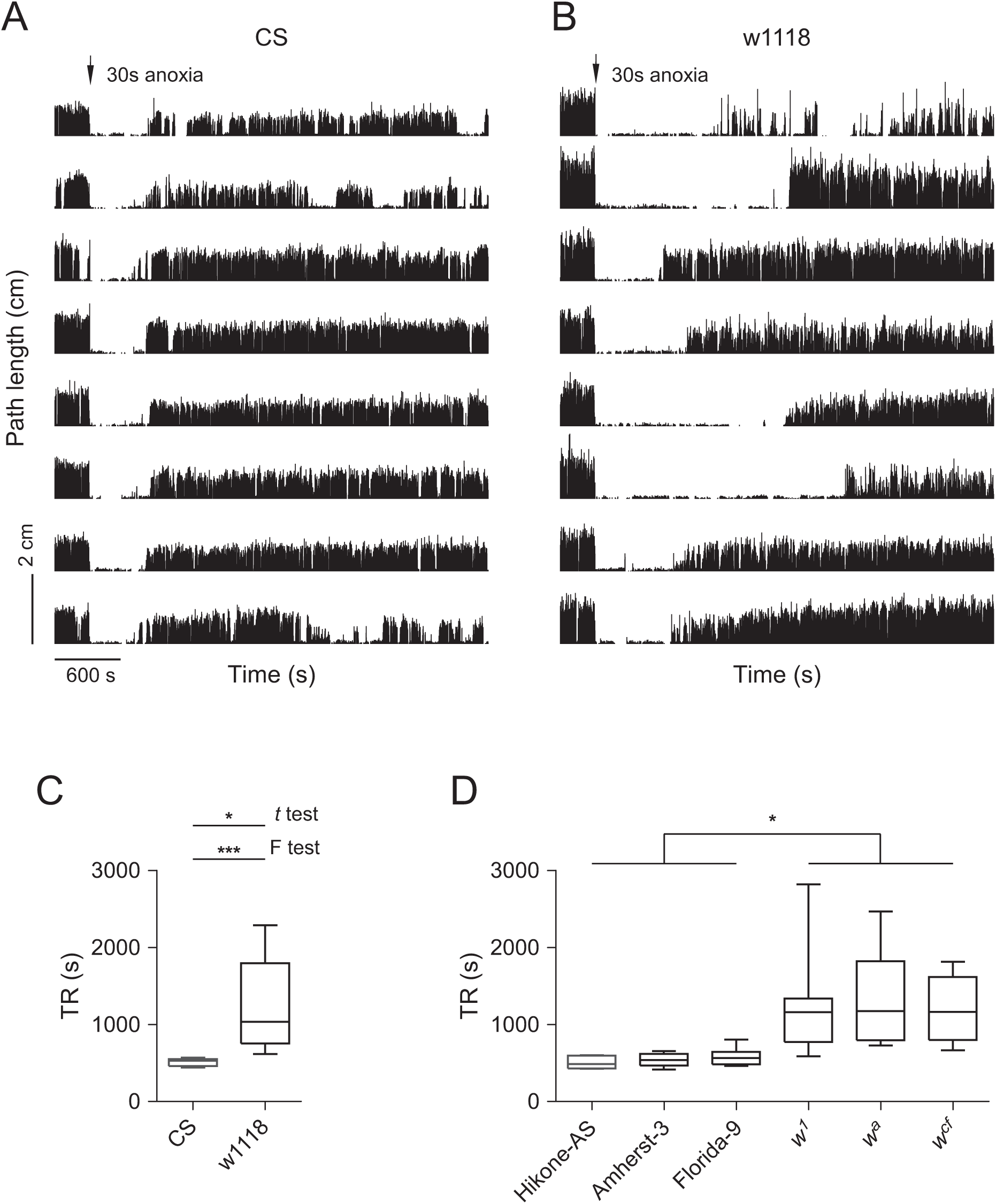
The *w* mutants display delayed locomotor recovery from anoxia. (A) Locomotor activity of CS flies in the circular arenas (1.27 cm diameter and 0.3 cm depth). Each trace represents 1h locomotion of a single fly. The path length per second vs time are plotted. A 30s anoxia (arrow head) is applied during 300 - 330 seconds. Flies are otherwise provided with normal air flow (at 2 L/min). (B) Locomotor activity of w1118 flies in the same arenas and subjected to the same treatment as CS. (C) Analysis of time to recovery (TR) in CS and w1118 flies. * *P* < 0.05 (t test), *** *P* < 0.001 (F test). (D) TR analysis in wild-types (Hikone-AS, Amherst-3 and Florida-9) and mutants (*w*^1^, *w*^a^ and *w*^cf^). * P < 0.05 by Kruskal-allis test.

### Mutants of *w* showed delayed locomotor recovery from anoxia

To determine whether the *w* mutation could be involved in the delayed locomotor recovery from anoxia, several additional wild-type strains including Hikone-AS, Amherst-3 and Florida-9 captured in different locations and dates, as well as the *w* mutants including *w^1^* (O’Hare *et al*. 1984), *w^a^* (Gehring and Paro 1980; Bingham and Judd 1981; Pirrotta and Bröckl 1984) and *w^cf^* (Mackenzie *et al*. 1999) were examined. The TRs in Hikone-AS (median 485.0 s, IQR 429.8 - 593.8 s, n = 8), Amherst-3 (median 538.5 s, IQR 466.5 - 621.0 s, n = 8) and Florida-9 (median 562.5 s, IQR 483.0 - 646.5 s, n = 8) were comparable with no significant difference. The TRs in *w^1^* (median 1162.0 s, IQR 772.5 - 1338.0 s, n = 12), *w^a^* (median 1173.0 s, IQR 795.0 - 1822.0 s, n = 15) and *w^cf^* (median 1166.0 s, IQR 797.0 - 1617.0 s, n = 8) were similar with no statistically significant difference. The *w* mutants displayed prolonged TRs compared with Hikone-AS (*P* < 0.05 for all three comparisons, Kruskal-Wallis test with Dunn’s Multiple Comparison), Amherst-3 (*P* < 0.05, Kruskal-Wallis test with Dunn’s Multiple Comparison), or Florida-9 (*P* < 0.05, Kruskal-Wallis test with Dunn’s Multiple Comparison) (Figure 1D). Thus, the *w* mutants showed delayed locomotor recovery from anoxia compared with wild-types.

### *w*^+^ allele was associated with fast locomotor recovery from anoxia

The w1118 strain carries isogenic X, second and third chromosomes and a null allele *w^1118^* on the X chromosome (Pirrotta and Bröckl 1984; Levis *et al*. 1985a). w1118 differs from CS in two main genetic components: the *w* allele and genetic background. The fast locomotor recovery from anoxia in wild-type might be associated with the *w*^+^ allele or the genetic background excluding the *w* locus. We next explored the genetic contribution of the *w*^+^ allele or genetic background to the timing of locomotor recovery.

By crossing male w1118 to female CS, and reciprocally, male CS to female w1118, two F1 male progenies were generated: *w*^+^/y (F1) and *w^1118^*/y (F1), which carried different X chromosomes and the same heterozygous genetic background on the second and third chromosomes. The median TR in *w*^+^/y (F1) flies was 424.0 s (IQR 411.5 - 428.0 s, n = 8), whereas the median TR in *w^1118^*/y (F1) flies was 662.5 s (IQR 613.0 - 786.5s, n = 8). The locomotor recovery in *w*^+^/y (F1) flies was earlier compared with *w^1118^*/y (F1) flies (*P* < 0.05, Mann-Whitney test) (Figure 2A).

**Figure 2.**
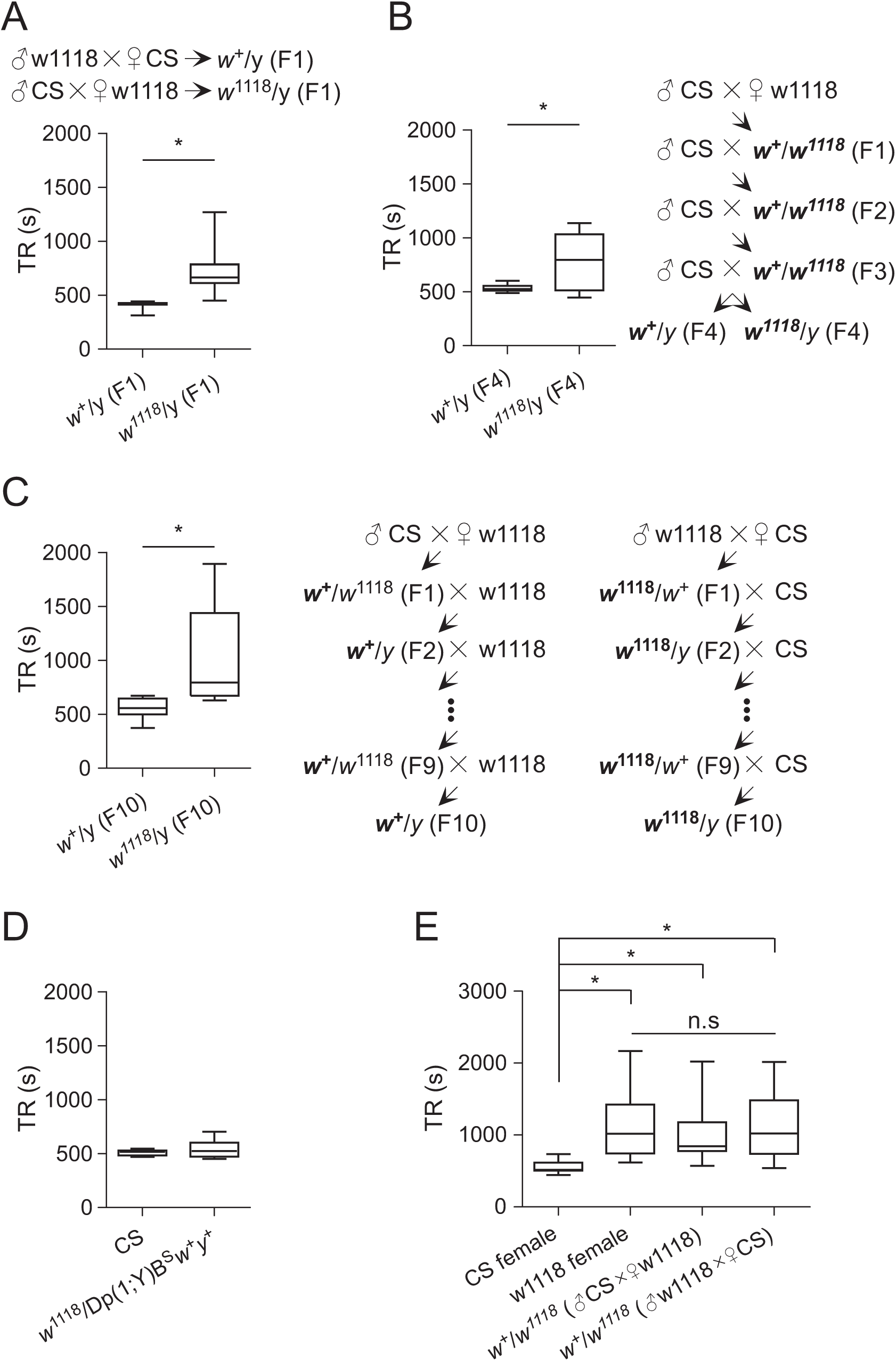
*w*^1118^ allele is associated with delayed locomotor recovery from anoxia. (A) TR analysis in the progeny flies from the cross between CS and w1118. The specific crossed are indicated. * *P* < 0.05 by Man-Whitney test. (B) TR analysis in flies generated by introgression (see Materials and Methods). * *P* < 0.05 by Man-Whitney test. (C) TR analysis in flies generated by serial backcrossing. * *P* < 0.05 by Man-Whitney test. (D) TR analysis in CS and flies carrying a *w*^+^ duplicated to Y chromosome. (E) TR analysis in these female flies: CS, w1118, *w*^+^/*w*^1118^ (progeny of male CS and female w1118) and *w*^+^/*w*^1118^ (progeny of male w1118 and female CS). * *P* < 0.05 by Kruskal-allis test; n.s, non-significant difference.

To examine the contribution of genetic background from wild-type, introgression (Kain *et al*. 2012) was performed to gradually replace isogenic second and third chromosomes in w1118 with wild-type counterparts. Briefly, by crossing male CS to female w1118, the heterozygous female progeny (F1) were collected and backcrossed to CS males. The backcross was conducted for three consecutive generations. The resulting *w*^+^/y (F4) and *w^1118^*/y (F4) flies contained synchronized and mostly wild-type genetic background. The median TRs were 525.5 s (IQR 509.0 - 554.5 s, n = 8) in *w*^+^/y (F4) flies, and 797.5 s (IQR 512.3 - 1032.0 s, n = 8) in *w^1118^*/y (F4) flies. The TR was shorter in *w*^+^/y (F4) flies than *w^1118^*/y (F4) flies (*P* < 0.05, *t* test with Welch’s correction) (Figure 2B). Hence, the fast locomotor recovery was tightly associated with *w*^+^-carrying flies.

To further examine the association between *w*^+^ allele and fast locomotor recovery, serial backcrossing was conducted to exchange *w* alleles between CS and w1118. Initially, male CS was crossed to female w1118, and reciprocally male w1118 was crossed to female CS. The heterozygous female progeny (F1) was backcrossed with w1118 or CS respectively. The backcrossing was performed for nine generations. The *w*^+^/y (F10) and *w^1118^*/y (F10) flies were collected and examined. The median TRs were 556.0 s (IQR 499.8 - 645.3 s, n = 8) in *w*^+^/y (F10), and 793.5 s (IQR 673.3 - 1439.0 s, n = 8) in *w^1118^*/y (F10). The TR was shorter in *w^1118^*/y (F10) flies that *w*^+^/y (F10) flies (*P* < 0.05, *t* test with Welch’s correction) (Figure 2C). Thus the *w*^+^ allele was strongly associated with fast locomotor recovery from anoxia.

### Flies with *w*^+^ duplicated to Y chromosome displayed fast locomotor recovery

To further support the association between *w* allele and the phenotype of locomotor recovery, we examined TR in flies carrying *w*^+^ duplicated to Y chromosome. The TR in *w*^1118^/Dp(1;Y)B^S^*w*^+^*y*+ (median 523.0 s, IQR 472.8 - 600.0 s) was similar to the TR in CS males (median 517.0 s, IQR 484.3 - 530.3 s) with no significant difference (Figure 2D). Thus, flies carrying *w*^+^ duplicated to the Y chromosome displayed fast locomotor recovery from anoxia.

### *w*^+^ allele was haplo-insufficient for fast locomotor recovery in female flies

We next addressed whether *w*^+^ allele was haplo-sufficient to promote fast locomotor recovery from anoxia in heterozygous female flies. The TR was examined in four different female flies: (1) CS, (2) w1118, (3) *w*^+^/*w^1118^* from the cross between male CS and female w1118, and (4) *w*^+^/*w^1118^* from the cross between male w1118 and female CS. The TR was at a median of 514.5 s (IQR 506.0 - 615.5s, n = 8) for CS females, 1018.0 s (IQR 745.5 - 1424.0 s, n = 8) for w1118 females, 846.0 s (IQR 776.8 - 1177.0 s, n = 8) for *w*^+^/*w^1118^* females (from male CS to female w1118), and 1022.0 s (IQR 737.8 - 1482.0s, n = 8) for *w*^+^/*w^1118^* females (from male w1118 to female CS). The TR in CS females was shorter than that in w1118 females (*P* < 0.05, ANOVA with Dunn’s multiple comparison), that in *w*^+^/*w^1118^* females (from male CS to female w1118) (*P* < 0.05, ANOVA with Dunn’s multiple comparison), and that in *w*^+^/*w^1118^* females (from male w1118 to female CS) (*P* < 0.05, ANOVA with Dunn’s multiple comparison). There was no difference of TR among w1118, *w*^+^/*w^1118^* (from male CS to female w1118) and *w*^+^/*w^1118^* (from male w1118 to female CS) (Figure 2E). These results indicated that homozygous *w*^+^ alleles were sufficient, whereas heterozygous *w*^+^ allele was insufficient to promote fast locomotor recovery from anoxia.

### Independent phenotypes between locomotor recovery from anoxia and eye pigmentation

The White protein carries and deposits red and brown pigments into the pigment cells of compound eyes, ocelli, Malpighian tubules and the testis sheath (Nolte 1950; Hazelrigg 1987). In addition to its classic role in pigmentation, the *w* allele is involved in several extra-retinal functions, inducing male-male courtship behavior (Zhang and Odenwald 1995), resistance to general volatile anesthetics (Campbell and Nash 2001), bio-amines transport (Borycz *et al*. 2008), positive phototatic personality (Kain *et al*. 2012) and place memory (Sitaraman *et al*. 2008). We looked for a possible association between phenotypes of locomotor recovery from anoxia and eye pigmentation. Flies carrying *cinnabar, brown (cn, bw)* double mutations in a *w*^+^ background were generated. These flies (*w*^+^; *cn,bw)* were white-eyed (Figure 3A) due to complete blockage of both pteridine and ommochrome pigment pathways (Waaler 1921; Ward 1923; Nolte 1954; Ghosh and Forrest 1967; Dreesen *et al*. 1988). There was no difference of TR between *w*^+^; *cn,bw* flies (median 579.0 s, IQR 467.8 - 674.8 s, n = 16) and CS (median 494.0 s, IQR 455.0 - 555.5 s, n = 16) (Figure 3B). Hence, the phenotype of eye pigment was dissociated with the timing of locomotor recovery.

**Figure 3.**
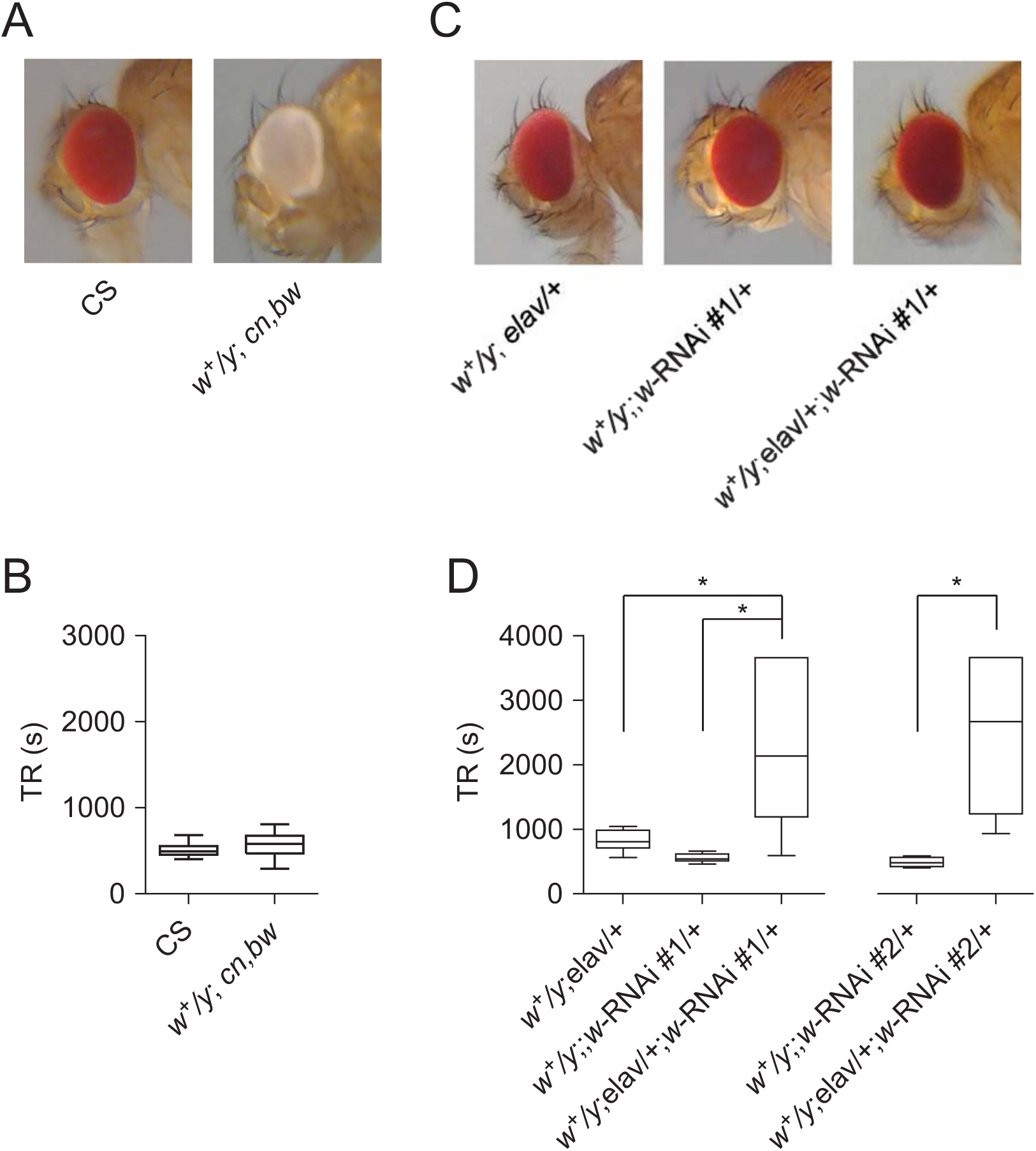
Independent phenotypes between locomotor recovery and eye pigmentation. (A) Eye color in CS and flies carrying *cinnabar* (*cn*) and *brown* (*bw*) double mutations (*w*^+^; *cn, bw*). (B) TR analysis in CS and *w*^+^; *cn, bw*. (C) Eye color in flies with *w*-RNAi targeted pan-neuronally to the central nervous system by elav-Gal4 and controls. (D) TR analysis in flies with *w*-RNAi knockdown and their controls. RNAi #1 and RNAi #2 indicate two independent RNAi lines targeting different sequences of *w* mRNA. * *P* < 0.05 by Kruskal-allis test.

To support the dissociation between locomotor recovery and pigmentation phenotype, RNAi knockdown of *w* was carried out. The eye color and the timing of locomotor recovery in flies with *w* knockdown were examined. The pan-neuronal driver elav-Gal4 (Armstrong *et al*. 2011; Dimitroff *et al*. 2012) was used for targeting RNAi of *w* to the central nervous system. There was no visible difference in eye color between *w*^+^; elav-Gal4/+; UAS-*w*-RNAi/+ flies and controls (*w*^+^; elav-Gal4/+ and *w*^+^;; UAS-*w*-RNAi/+) (Figure 3C), indicating that *w* knockdown in the central nervous system had no visible effect on eye pigmentation. The TR of *w*^+^; elav-Gal4/+; UAS-*w*-RNAi/+ males was, however, severely delayed compared with controls (Figure 3D). The RNAi-induced delay of locomotor recovery was confirmed by using two independent RNAi lines targeting different sequences of the *w* transcript. These results clearly display an independence between phenotypes of locomotor recovery from anoxia and eye pigmentation.

### mini-*white* (m*w*^+^) was insufficient to promote fast locomotor recovery

The m*w*^+^ contains reduced 5’ and 3’ regulatory sequences and a greatly shortened first intron with a deletion of ~3kb HindIII - Xbal fragment (Pirrotta 1988). Depending on its location in the genome, m*w*^+^ displays a more or less cell-autonomous expression pattern and is controlled by cis-acting regulatory sequences (Hazelrigg *et al*. 1984; Hazelrigg 1987). In the *Drosophila* community most transgenic lines carry the m*w*^+^ marker. Because both *w*^+^ and m*w*^+^ encode the same protein, we examined the contribution of inserted m*w*^+^ to the timing of locomotor recovery. Importantly, the contribution of m*w*^+^, if any, needs to be differentiated from the *w*^+^ allele if the m*w*^+^-carrying Gal4/UAS expression system (Brand and Perrimon 1993) is introduced into the wild-type genetic background. There were several criteria for these tests: (1) UAS but not Gal4 lines were selected in order to avoid potential modulation by ectopic Gal4 expression (Kramer and Staveley 2003; Rezával *et al*. 2007); (2) UAS lines with m*w*^+^-carrying P elements inserted into fixed recombination sites at attP2 or attP40 were preferentially considered in order to minimize the position effect (Nolte 1950; Hazelrigg *et al*. 1984); (3) UAS flies carrying multiple copies of m*w*^+^ were generated to compensate for possibly reduced expression from a single m*w*^+^.

We first measured TR in flies carrying P element (pJFRC2) at attP40 on the second chromosome or P element (pJFRC2, pJFRC5 or pJFRC7) at attP2 on the third chromosome (Pfeiffer *et al*. 2010). All the selected UAS lines carry the m*w*^+^ marker. They were backcrossed into w1118 for 10 generations before experiments by following previously described procedures (Garfinkel *et al*. 2004). Flies containing multiple copies of P elements were generated by genetic manipulation (Greenspan 2004). We found: (1) TR of each heterozygous UAS line carrying a single m*w*^+^ was comparable with that of w1118 with non-significant difference (non-parametric ANOVA with Dunn’s multiple comparison) (Figure 4A); (2) TR of each homozygous UAS line or the combination of UAS containing two m*w*^+^ was similar to that of w1118 with no statistically significant difference (non-parametric ANOVA with Dunn’s multiple comparison) (Figure 4B); (3) TR of each line carrying four copies of m*w*^+^ was statistically the same as that in w1118 (non-parametric ANOVA with Dunn’s multiple comparison) (Figure 4C). Therefore, up to four copies of m*w*^+^ at attP40 or attP2 in the w1118 background were insufficient to promote fast locomotor recovery.

**Figure 4.**
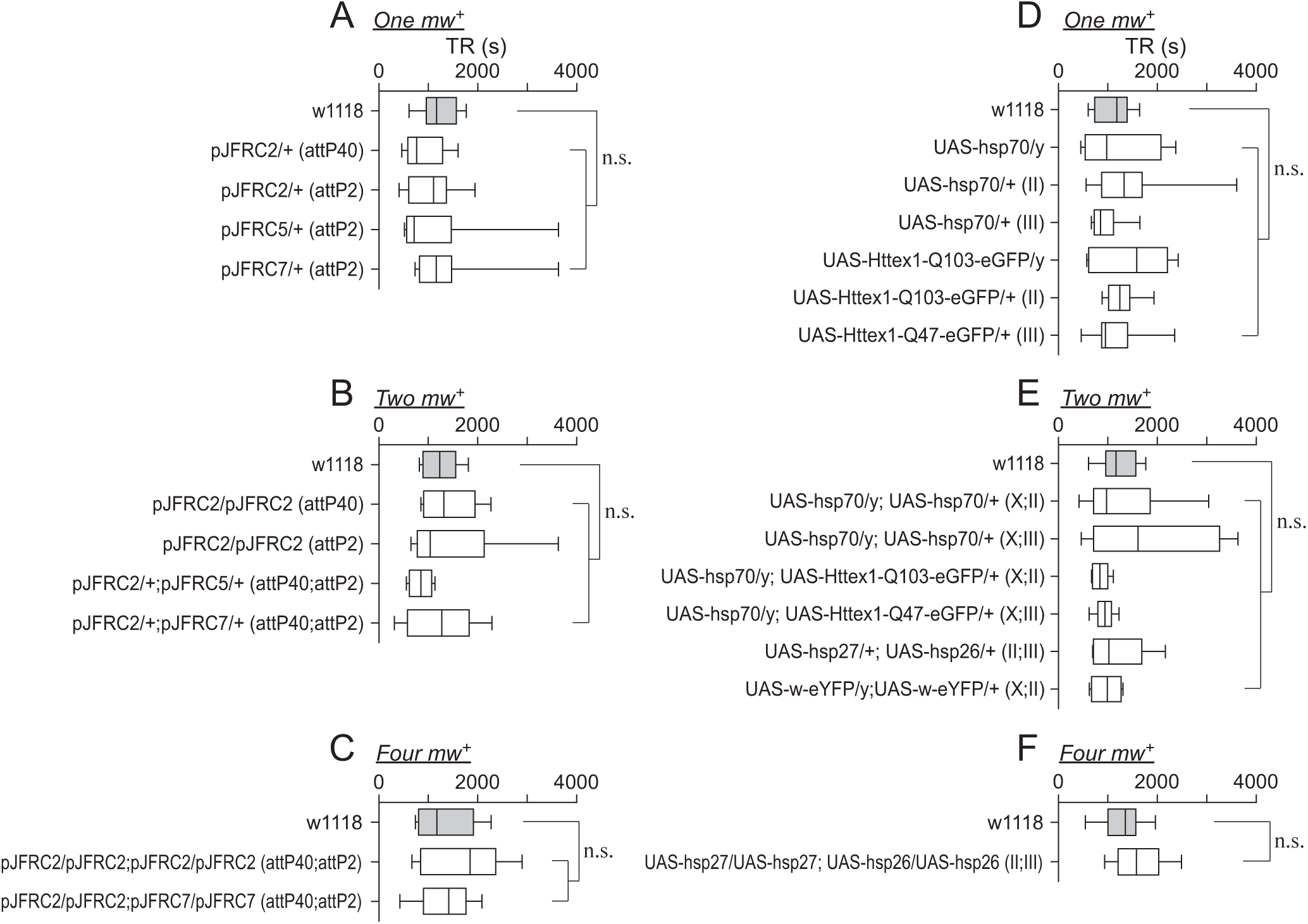
The m*w*^+^ is insufficient to promote fast locomotor recovery. (A, B, C) TR analysis for flies carrying 1-4 copies of m*w*^+^ at attP40 or attP2 docking site by site-specific recombination. pJFRC2, pJFRC5 and pJFRC7 are the UAS constructs (Pfeiffer *et al*. 2010). The specific location of m*w*^+^ for each fly line is indicated in the parenthesis. n.s, non-significant difference. (D, E, F) TR analysis for flies carrying 1-4 copies of m*w*^+^ at nearly random locations. The chromosome location for each line is indicated. n.s, non-significant difference. Notes: Flies carrying pJFRC2-7 were backcrossed into w1118 for 10 generations before the tests. UAS-hsp70 flies are constructed in the lab and reported previously (Xiao *et al*. 2007). The other fly lines and sources are: UAS-Httex1-Qn-eGFP (n = 47 or 103) (Zhang *et al*. 2010); UAS-hsp26 and UAS-hsp27 (Wang *et al*. 2004); UAS-w-eYFP (Evans *et al*. 2008).

Next we examined TR in UAS flies with nearly random locations of m*w*^+^ on X, second or third chromosome. We chose several UAS-hsp70 lines with pINDY5-hsp70 inserted into different chromosomes (Xiao *et al*. 2007), fly lines carrying UAS-Httex1-Qn-eGFP (n = 47 or 103) in different chromosomes (Zhang *et al*. 2010), two different lines of UAS-*w*-eYFP (Evans *et al*. 2008), and a few other UAS flies in our collection. All of the selected UAS flies were generated in the w1118 background, contained m*w*^+^ marker, and displayed no apparent developmental or morphological abnormalities. TR in each of the fly lines with m*w*^+^ copy number ranging from one (Figure 4D), two (Figure 4E) to four (Figure 4F) was similar with that in w1118 with non-significant difference (non-parametric ANOVA with Dunn’s multiple comparison). Thus, up to four copies of m*w*^+^ with nearly random insertions in the genome were insufficient to confer fast locomotor recovery. Notably, the UAS lines with a P element on X chromosome (UAS-hsp70/y and UAS-Httex1-Q103-eGFP/y) displayed TR similar with that in w1118, indicating that expression of m*w*^+^ with potential dosage compensation (Pirrotta and Bröckl 1984; Qian and Pirrotta 1995) was unable to promote fast locomotor recovery.

### Knockdown of *w* in putative subsets of serotonin neurons was sufficient to delay locomotor recovery

We demonstrated above that RNAi knockdown of *w* pan-neuronally using elav-Gal4 severely delayed locomotor recovery (see Figure 3D). These data suggest that White is expressed in CNS neurons. We further examined the possible expression of White in glial cells. RNAi knockdown of *w* throughout glia using repo-Gal4 (Sepp *et al*. 2001) with two independent RNAi lines targeting different sequences of *w* transcript had no effect on TR (Figure 5A). These data indicate no or minor expression of White in glial cells.

**Figure 5.**
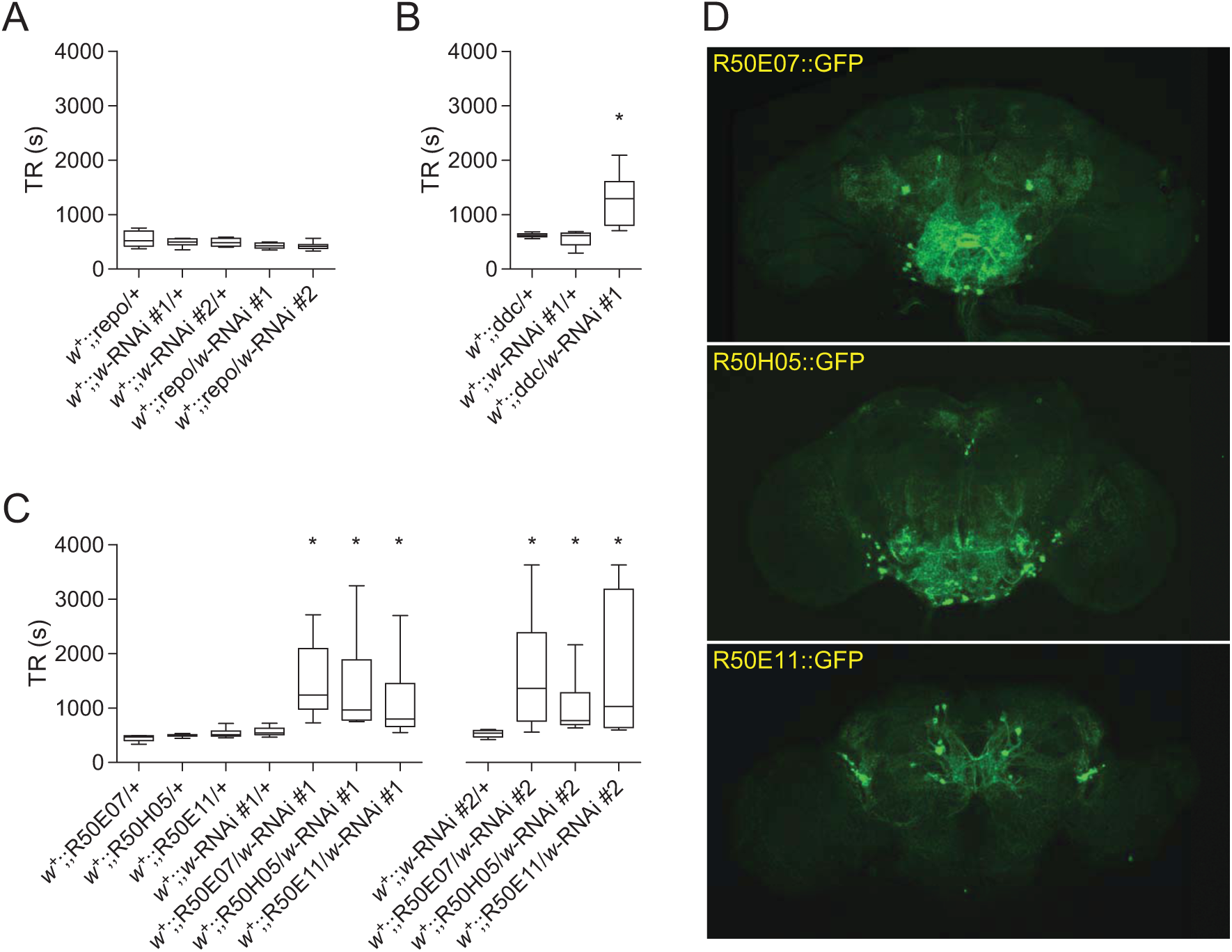
RNAi knockdown of *w* in subsets of putative serotonin neurons is sufficient to delay locomotor recovery. (A) TR analysis for flies with *w*-RNAi knockdown in pan-glial cells by repo-Gal4 (Ref). TR was unaffected by w-RNAi in pan-glial cells. (B) TR analysis for flies with *w*-RNAi knockdown by ddc-Gal4. Ddc-Gal4 is specific to dopaminergic and serotonergic neurons (Haywood and Staveley 2004). (C) TR analysis for flies with *w*-RNAi knockdown by R50E07-, R50H05- and R50E11-Gal4. These Gal4 lines use the enhancers originated respectively from three major introns from *SerT* (serotonin transporter gene) (Pfeiffer *et al*. 2008). Two independent RNAi lines (#1 and #2) were used for the tests. * *P* < 0.05 by Kruskal-allis test. (D) Expression patterns of R50E07-, R50H05- and R50E11-Gal4. Each driver targets 20 - 50 neurons in the central brain. The GFP denotes 20×UAS-IVS-mCD8::GFP fly line (#32194, Bloomington stock center).

White is likely expressed in dopaminergic and serotonergic neurons, because a co-enzyme tetrahydrobiopterin is required for biosynthesis of both dopamine and serotonin, and tetrahydrobiopterin is an important co-product in the red pigment pathway that requires White (Ferré *et al*. 1986). We next performed RNAi knockdown of *w* using ddc-Gal4 (specific to dopaminergic and serotonergic neurons) and examined the effect on the timing of locomotor recovery. TR in *w*^+^;; ddc/*w*-RNAi flies (median 1298.0 s, IQR 805.0 - 1608.0 s, n = 8) was clearly prolonged compared with *w*^+^;; ddc/+ (median 618.0 s, IQR 593.0 - 647.5s, n = 8) or *w*^+^;; *w*-RNAi/+ flies (median 616.0 s, IQR 447.3 - 662.0s, n = 8) (*P* < 0.05, non-parametric ANOVA with Dunn’s multiple comparisons) (Figure 5B). Results support White expression in dopaminergic and serotonergic neurons and that knockdown of *w* in these neurons delayed locomotor recovery from anoxia.

Serotonin synthesis begins with hydroxylation of tryptophan, which is also the initial substrate for production of brown pigment xanthommatin, a process requiring White (Sullivan and Sullivan 1975). Serotonin content is greatly reduced in the mutant strain w1118 (Borycz *et al*. 2008; Sitaraman *et al*. 2008). We then examined whether knockdown of *w* in putative serotonin neurons was sufficient to delay locomotor recovery. Three Janelia Gal4 lines, R50E07-, R50H05- and R50E11-Gal4, using respectively one of the three large intron regions of the serotonin transporter gene (*SerT*) as enhancers, potentially divide the serotonin population into smaller subsets (Pfeiffer *et al*. 2008). RNAi knockdown by each of the three Gal4 lines severely delayed locomotor recovery compared with the controls. The delayed locomotor recovery was observed consistently by using two independent RNAi lines (Figure 5C). The expression patterns of R50E07-, R50H05- and R50E11-Gal4 are illustrated in Figure 5D. Each Gal4 line labeled 20 - 50 neurons in the central brain, which were smaller populations than the entire serotonergic family of around 100 neurons (Vallés and White 1988; Giang *et al*. 2011; Huser *et al*. 2012). The labeling was highly consistent with reported expression pattern (Pfeiffer *et al*. 2008). Therefore, knockdown of *w* in each of the putative serotonin subsets was sufficient to cause delayed locomotor recovery.

## Discussion

We report here a novel function of *w* that is tightly associated with the timing of locomotor recovery from anoxia. Down-regulation of White, either by mutation or RNAi knockdown pan-neuronally, delayed locomotor recovery. Genetic analysis through serial backcrossing together with simple crossing and introgression confirmed the strong association between *w* and locomotor recovery phenotype. We further demonstrate the independence of the phenotypes between locomotor recovery and eye pigmentation, although they both are associated with *w*. Moreover, we show that *w*^+^ is haplo-insufficient for fast locomotor recovery in female flies, and that up to four copies of m*w*^+^ are insufficient to promote fast locomotor recovery. Nevertheless, flies carrying *w*^+^ duplicated to Y chromosome display wild-type like fast locomotor recovery. Finally, we show that RNAi knockdown of *w* in subsets of putative serotonin neurons is sufficient to delay locomotor recovery from anoxia.

### The expression pattern of White protein

White protein is expressed in vesicular but not cytoplasmic membranes in principle cells of Malpighian tubules (Evans *et al*. 2008). Expression of White can also be detected in the granules of pigment cells in the lamina and in photoreceptors (Mackenzie *et al*. 2000; Borycz *et al*. 2008). However, a convincing expression pattern in the CNS is still lacking. We provide supportive evidence for the expression patterns of White protein in the CNS. White is likely expressed in neurons but not glial cells. This is supported by the observations that the delayed locomotor recovery can be induced by RNAi knockdown of *w* pan-neuronally or in subsets of putative serotonin neurons but not in glia.

The level of White in CNS neurons would be extremely low based on the report of unsuccessful immunostaining (Borycz *et al*. 2008). Meanwhile, the expression is tightly controlled by the adjacent regulatory array of *w* gene (Hazelrigg *et al*. 1984; Hazelrigg 1987). Insertion of m*w*^+^ results in abnormal sexual performance in male flies (Zhang and Odenwald 1995) and partial recovery of eye color ranging from light yellow to dark red, suggesting that m*w*^+^ either causes behavioral abnormality, or is insufficient to rescue the loss-of-*w* function. Consistently, I observed that the insertion of m*w*^+^ into a mutant background was unable to promote fast locomotor recovery that was typically seen in wild-type flies. It is argued that abnormal expression as well as intracellular mis-location of White in normally expressing cells is associated with male-male courtship behavior (Anaka *et al*. 2008; Krstic *et al*. 2013). Hence, *w*^+^ and its expression might represent a highly optimized system in eye pigmentation and substrates transport. Complete lack or down-regulation of White could break down such an optimized system and lead to a prolonged time for restoration of vesicular content during locomotor recovery.

### Multiple roles of White protein

For almost a century White has been known to control eye pigmentation pathways. It was not until 1995 that White was proposed to be associated with male-male courtship behavior (Zhang and Odenwald 1995). Recently, more neural but non-retinal functions of White have been revealed (Campbell and Nash 2001; Borycz *et al*. 2008; Krstic *et al*. 2013). In this study, we demonstrate a clear independence of phenotypes between eye pigmentation and the timing of locomotor recovery. We consider below several factors including temporal expression pattern, cell-specificity and reusability of White-containing granules/vesicles that might explain the potential multiple roles for White.

In the eyes of wild-type flies, White-containing granules carry and deposit two independent eye pigments to the same granules in the pigment cells. The brown pigment is deposited first with a peak being reached only about 1 day after emergence; then the red pigment is deposited in the same granules for another 4 or 5 days (Nolte 1950). After that, White-containing granules might complete the deposition of pigments. The one-way centrifugal movement of pigment granules and the unchangeable color pattern throughout the life in the mutant *w^m4^* (Nolte 1950) imply that White-containing granules are either not able to return to the cell soma or do so at a very low rate for renewed pigment transport. In the CNS, however, the function of White-containing vesicles could persist. The observation that in flies up to 9 days old down-regulation of White in neurons delays locomotor recovery supports the idea that White continues to function in the adult for more than 4 or 5 days. Also, up to 10 day-old male flies with m*w*^+^ still display male-male courtship behavior, which is believed to be controlled by ~2000 neurons in the CNS (Manoli *et al*. 2005; Hampel *et al*. 2011), indicating persistent function of White in CNS. Thus, the multiple roles of White are supported by temporal expression patterns from difference tissues.

CNS-located serotonin neurons provide a tissue basis for additional roles for White other than in controlling eye color in pigment cells. These neurons release serotonin (5HT), which acts as a neuromodulator involved in locomotor behavior, circadian entrainment and place memory (Yuan *et al*. 2005; Sitaraman *et al*. 2008; Nall and Sehgal 2014; Okusawa *et al*. 2014; Silva *et al*. 2014). Recovery from anoxia includes the restoration of 5HT vesicular stores that would be excessively released during anoxic coma (Hansen 1985). Normal expression of White increases vesicular content of 5HT (Borycz *et al*. 2008; Sitaraman *et al*. 2008), suggesting the presence of White in serotonin neurons. Here we show that RNAi knockdown of *w* in dopaminergic and serotonergic neurons, or putative subsets of serotonin neurons delays locomotor recovery, providing further evidence that White is expressed in the CNS and possesses potentially non-retinal function.

The molecular basis for multiple roles of White is perhaps related to its function of non-specific substrate transport. White, together with its transporting partners, accumulates a variety of small molecules into granules or vesicles (Anaka *et al*. 2008). The formation of vesicular stores preserves a sufficient amount of neurotransmitters or second messengers and improves signaling efficacy. Down-regulation of White would increase the chances of exposing transmitters and other substrates to cytosolic enzymatic environments and would result in excessive hydrolysis (Evans *et al*. 2008). This is supported by the observation that White down-regulation shifts the timing of locomotor recovery from consistently fast to remarkably slow with large variance.

### Insufficient rescue of delayed locomotor recovery by m*w*^+^ and its implication

Intronic sequences of *w* might carry tissue-specificity or unknown regulatory information (Pirrotta and Bröckl 1984; Hazelrigg 1987; Qian and Pirrotta 1995). Removal of a large intronic fragment of the *w* gene could impair transcriptional regulation. Similarly, the relocation of m*w*^+^ in the genome could pick up additional regulatory information. If the *w*^+^ system were extremely optimized, the insertion of m*w*^+^ would result in either no or a very limited effect on the rescue of locomotor recovery. Additionally, the fact that heterozygous (*w*^+^/*w^1118^*) female flies showed no difference from mutant w1118 in the timing of locomotor recovery implies a haplo-insufficient function of *w*^+^ in the females. Although in male flies dosage compensation is another critical mechanism in increasing the transcription of a single *w* allele (Hazelrigg 1987; Qian and Pirrotta 1995), the relocation of m*w*^+^ might result in the abnormal regulation of dosage compensation.

Although mutants of *w* and flies with RNAi knockdown of *w* in putative serotonin subsets demonstrate the same outcome – delayed locomotor recovery from anoxia, the genetic underpinnings are clearly different. Down-regulation of White through an RNAi approach in restricted cell populations is sufficient to delay locomotor recovery, and down-regulation in different subsets yields a uniform delay. Each of the putative serotonin subsets consists of fewer than 100 neurons in CNS. The uniform delay induced in such a small population indicates a critical role for White in highly specified or differentiated neurons. This could explain the insufficiency to rescue the timing of locomotor recovery in w1118 flies by genetic expression of m*w*^+^. Most likely, expression levels and tissue patterns must be fully recovered to generate the wild-type phenotype. However, m*w*^+^ expression is subjected to strong regulation from adjacent cis-acting but non-native elements (Hazelrigg *et al*. 1984; Levis *et al*. 1985b; Levis *et al*. 1985a). Taking into account a loss of a large intronic sequence that might carry essential regulatory elements for normal expression, and that dosage compensation must also be normal to generate the wild-type phenotype, rescuing the locomotor recovery phenotype in the w1118 mutant might not be possible by genetic manipulation of m*w*^+^.

Notwithstanding the insufficient rescue of locomotor recovery phenotype by m*w*^+^, flies carrying *w*^+^ duplicated to the Y chromosome (Brosseau *et al*. 1961) display wild-type like, fast locomotor recovery from anoxia. Such a complete rescue supports the association between the *w* gene and locomotor recovery phenotype.

Future work should focus on the molecular mechanisms underlying the *w*-associated fast locomotor recovery from anoxia. Particularly, identifying the specific transporting substrates that are responsible for the timing of locomotor recovery would be promising.

